# Cell-type specific astrocyte activation is driven by cortical top-down modulation

**DOI:** 10.64898/2026.03.08.710364

**Authors:** Antonia Beiersdorfer, Kristina Losse, Jennifer Bostel, Janina Sophie Popp, Natalie Rotermund, Kristina Schulz, Damian Droste, Christine E. Gee, Daniela Hirnet, Christian Lohr

**Author notes:** Corresponding authors: Christian Lohr, and Antonia Beiersdorfer. Equal contribution (Antonia Beiersdorfer, Kristina Losse).

## Abstract

Cortical projections to cortical and subcortical targets provide top-down modulation that shapes neuronal performance, including gain control and excitation–inhibition balance. However, the contribution of astrocytes to this process remains poorly understood. In the olfactory bulb, the first relay station of odor information processing, bottom-up input is transmitted from olfactory sensory neurons to mitral/tufted (M/T) cells, which project to the olfactory cortex. Context- and state-dependent top-down modulation arises from feedback projections originating in the anterior piriform cortex (aPC) that target granule cells (GCs). We examined how astrocytes respond to bottom-up and top-down neuronal activity using confocal Ca²⁺ imaging, cell-type-specific optogenetics, electrical stimulation, and single-cell electrophysiology. We found that Ca²⁺ signals in astrocytes are selectively triggered by action potential-dependent ATP release from GCs while M/T cells failed to elicit significant astrocytic responses. Although synaptic input from M/T cells depolarized GCs, it was insufficient to induce action potential firing and subsequent astrocyte activation. By contrast, glutamatergic top-down input from the aPC evoked sustained GC firing, leading to ATP-dependent Ca²⁺ signaling in astrocytes. Our results reveal an unappreciated level of complexity in neuron–astrocyte communication, highlighting its cell-type specificity as well as its context- and state-dependence.

## Introduction

One of the most recent achievements in the evolution of the mammalian brain is the addition of the neocortex to pre-existing brain structures (Sherman & Usrey, 2021). Neocortical regions modulate most of our actions through “top-down” projections to subcortical areas such as the superior colliculus, the basal ganglia, and the thalamus (Cruz et al., 2023). For example, several cortical regions influence sensorimotor integration and attention both through direct glutamatergic projections and indirectly via loops involving the basal ganglia. Furthermore, functional coupling between the prefrontal cortex and the ventral tegmental area is critical for reward-related processing(Wu et al., 2013). Another example demonstrates that fear behavior is strongly regulated by top-down projections from the medial prefrontal cortex to the amygdala(Mack et al., 2023). Such cortical top-down modulation of neuronal activity not only occurs between cortical and subcortical regions, but also among cortical territories, in particular from higher to lower hierarchy cortical structures (Felleman & Van Essen, 1991; Wang et al., 2020). Cortico-cortical top-down modulation affects prediction and attention and tunes excitation-inhibition balance as well as gain in sensory cortices (Batista-Brito et al., 2018; Choi et al., 2018; Miller & Buschman, 2013). In the olfactory system, e.g., sparse and odor-specific feedback from pyramidal neurons of the anterior piriform cortex (aPC) projecting to the olfactory bulbs (OBs) results in release of glutamate that excites OB granule cells (GCs) (Boyd et al., 2015; Boyd et al., 2012; Li et al., 2020). GABA released from GCs upon feedback excitation from the piriform cortex mainly affects mitral cells (MCs) but not tufted cells (TCs), which are the two principal (output) neurons of the OB (Otazu et al., 2015). This top-down modulation is involved in behavioral responses such as recognition and valence of novel social odor cues as well as reward contingency (Hernandez et al., 2025; Zhou et al., 2025).

While the role of neurons in cortico-cortical and cortico-subcortical top-down modulation is well described, it is unknown how astrocytes are involved in this process. In addition to their homeostatic functions, astrocytes release gliotransmitters acting on neuronal receptors, thereby tuning neuronal activity, contributing to synaptic plasticity and shaping behavior which renders them ideal to mediate neuronal modulation evoked by top-down feedback inputs from the cortex (Durkee et al., 2021; Lyon & Allen, 2021; Mazaud et al., 2021). Most astrocytic functions are controlled or modulated by cytosolic Ca^2+^ signals (Lim et al., 2021). Ca^2+^ signaling in astrocytes can be evoked by the activation of metabotropic as well as ionotropic receptors as a result of neurotransmitter release from neurons (Ahmadpour et al., 2021; Caudal et al., 2020; Okubo & Iino, 2020). Among these neurotransmitters are glutamate, GABA, acetylcholine, adenosine-triphosphate (ATP), norepinephrine, and dopamine, which increase Ca^2+^ in astrocytes in many brain regions such as the cortex, hippocampus and cerebellum (de Melo Reis et al., 2020; Lim et al., 2021). Because astrocytes respond to a multitude of neurotransmitters, it was hypothesized that they integrate the activity of the entire neuronal population within the reach of the astrocytic cell processes with little or no neuronal cell-type specificity, which contradicts a specific role in mediating defined tasks in neuronal information processing such as cortical top-down modulation.

Astrocytes in the OB glomerular layer, in which incoming olfactory information is transmitted to mitral/tufted (M/T) cells and processed by interaction with local interneurons, respond to glutamate and ATP released from axon terminals of olfactory sensory neurons (Doengi et al., 2008; Thyssen et al., 2010). The output signal of M/T cells is then shaped in a deeper layer, the external plexiform layer (EPL), that is characterized by a high density of reciprocal dendrodendritic synapses between M/T cells and interneurons, the vast majority provided by axonless GCs (Bartel et al., 2015; Burton, 2017). These synapses mediate recurrent dendrodendritic inhibition; M/T cells release glutamate that excites GCs, which in turn release GABA to feedback on the M/T cells. How astrocytes are embedded in this neuronal microcircuit has not been studied so far and we asked the question whether astrocytes in the EPL distinguish between bottom-up neuronal signaling carried by M/T cell activity versus GC activity evoked by cortical top-down inputs.

To address neuron-astrocyte communication in the EPL, we used GCaMP6s-expressing astrocytes in acute brain slices of the OB to visualize astrocytic Ca^2+^ signaling. We established stimulation protocols to excite different neuron populations that form dendrodendritic synapses in the EPL. Electrical and optogenetic stimulation of GCs, but not M/T cells, resulted in large Ca^2+^ transients in astrocytes mediated by P2Y1 receptors. Surprisingly, astrocytes failed to respond to granule cell activity when GCs were synaptically excited by M/T cell stimulation, but responded to granule cell activity when GCs were excited by glutamatergic feedback projections from the aPC. Our results show that astrocytes can distinguish between different neuron populations and participate in cortical top-down modulation, making neuron-astrocyte transmission cell type-specific and state-dependent.

## Results

### Neuronal GCaMP expression in Pcdh21-Cre x GCaMP6s^flox^ mice

To visualize M/T cell activation and to establish a stimulation protocol that selectively activates M/T cells in the EPL of the OB, we took advantage of Pcdh21-Cre x GCaMP6s^flox^ mice, selectively driving the expression of the genetically encoded calcium indicator GCaMP6s in neurons of the OB (Fig. 1a). The Pcdh21 promoter encodes for a nonclassical cadherin that is predominantly expressed in M/T cells of the OB (Nagai et al., 2005; Wachowiak et al., 2013). We analyzed GCaMP6s expression in Pcdh21-Cre x GCaMP6s^flox^ mice in the OB via immunohistochemistry using anti-GFP antibodies (Fig. 1b-e). GCaMP6-positive MCs can easily be identified by their large cell bodies (> 20 µm), lined up next to each other in the mitral cell layer (ML) and projecting a primary apical dendrite towards the GL and secondary lateral dendrites horizontally in the EPL (Fig.1e, arrow 1). In addition, TCs located at the boundary between GL and EPL showed GCaMP6s expression (Fig.1e, arrow 2). This GCaMP6s expression pattern was found in 73 % of the analyzed mice, which shows that our mouse model is well suited to study Ca^2+^ signals in OB principal neurons (M/T cells). To visualize neuronal activity and Ca^2+^-dependent neurotransmitter release by M/T cells, we prepared acute OB slices and electrically stimulated different loci of the OB (Fig. 1b-c). M/T cells were directly stimulated using a stimulation pipette in the IPL (Fig. 1c), where the axons of M/T cells converge to form the lateral olfactory tract (LOT), in the presence of glutamatergic and GABAergic inhibitors. Stimulation in the IPL induced Ca^2+^ transients in M/T cells (Fig 1f, h). No significant difference was found in the amplitude of Ca^2+^ transients in MCs or and TCs nor between cell somata, apical dendrites and lateral dendrites (Fig. s1). When we instead stimulated the M/T dendritic tuft in the GL (Fig. 1c), Ca^2+^ transients were evoked in 78.3% of the MCs that had responded to IPL stimulation before (Fig. 1g-i).

**Figure 1:**
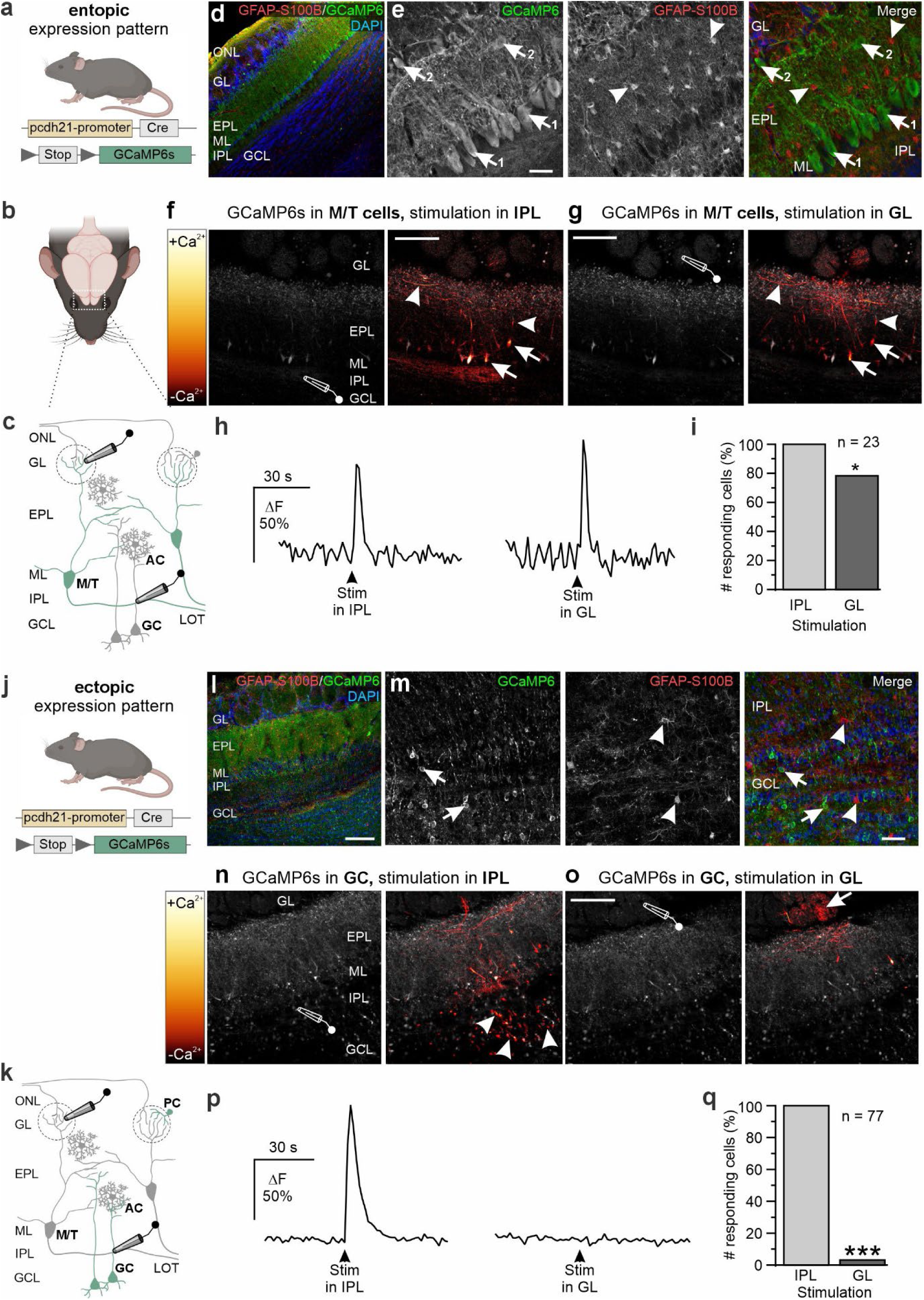
Electrical stimulation in different layers of the OB drives the activity of distinct populations of OB neurons. **(a)** Mouse model entopically expressing GCaMP6s controlled by the pcdh21-promoter. **(b)** Location of OBs. **(c)** Structure of the OB. Axons of olfactory sensory neurons enter the olfactory nerve layer (ONL) and synapse on mitral cells (MC) and tufted cells (TC) in the glomerular layer (GL). Dendrites of MCs extend through the external plexiform layer (EPL). Somata of MCs are located in the mitral cell layer (ML), their axons proceeding through the internal plexiform layer (IPL) building the lateral olfactory tract (LOT). Granule cells (GCs) are located the granular cell layer (GCL). Astrocytes (AC) are located throughout all layers of the OB. M/T cells expressing GCaMP6s are indicated in green. Stimulation pipettes indicated in the GL and IPL. **(d)** Entopic GCaMP6 expression (green). Astrocytes were S100b/GFAP immunostained (red), nuclei with Dapi (blue). Scale: 100 µm. **(e)** GCaMP6s was expressed by MCs (arrows 1) and tufted cells (arrows 2), but not astrocytes (arrowheads). Scale: 20 µm. **(f)** Image sequence before (left) and after (right) electrical stimulation in the IPL with M/T cells expressing GCaMP6s. Scale: 100 µm. **(g)** Electrical stimulation in the GL. Scale: 100 µm. **(h)** Ca^2+^ transients in mitral cells upon stimulation in the IPL and GL. **(i)** Fraction of MCs responding to stimulation in IPL and GL. **(j)** Expression of GCaMP6s controlled by the pcdh21-promoter. **(k)** Structure of the OB. GCs and periglomerular cells expressing GCaMP6s are indicated in green. **(l)** Ectopic GCaMP6 expression (green) in the OB of Pcdh21-Cre x GCaMP6s^flox^ mice. Scale: 100 µm. **(m)** GCs were GCaMP6s-positive (arrows), GFAP/S100b-immunopositive astrocytes did not express GCaMP6s (arrowheads). Scale: 20 µm. **(n)** Image sequence before (left) and after (right) electrical stimulation in the IPL with GCaMP6s in GCs (arrowheads). Scale: 100 µm. **(o)** Image sequence before (left) and after (right) electrical stimulation in the GL. No GCs but few dendritic structures belonging to the local network of the stimulated glomerulus responded to electrical stimulation (arrow). Scale: 100 µm. **(p)** Ca^2+^ transients in granule cells upon stimulation in the IPL and GL. **(q)** Fraction of GCs responding to stimulation in the IPL and GL. All experiments were performed in synaptic isolation: AMPA receptor antagonist: NBQX 20 µM; NMDA receptor antagonist: D-APV 100 µM; mGluR1 receptor antagonist: CPCCOEt 100 µM; GABA_A_ receptor antagonist: Gabazine 10 µM; GABA_B_ receptor antagonist: CGP55845 10 µM to ensure isolation of direct responses to electrical stimulation. *, p < 0.05; ***, p < 0.001.

In addition to the intended expression pattern, 27% of the Pcdh21-Cre x GCaMP6s^flox^ mice had a germ line mutation resulting in GCaMP6s expression in interneuron populations of the OB and no expression in M/T cells (Fig. 1j, l-m). In these mice, we found dominant GCaMP6s expression in OB GCs, identified by their abundance and localization in the granule cell layer (GCL) and basal ML, their small and densely packed somata and the projection of one apical dendrite into the EPL (Fig. 1l-m). In addition, a few GCaMP6s-positive juxtaglomerular interneurons were located in the GL. We termed GCaMP6s expression in interneurons “ectopic”, while we refer to GCaMP6s expression in M/T cells as “entopic”. No colocalization of GCaMP6s and GFAP/S100b was detected in both variants of Pcdh21-Cre x GCaMP6s^flox^ mice, indicating lack of GCaMP6s expression in astrocytes (Fig. 1e, m). We next analyzed Ca^2+^ transients in GCs in response to stimulation in the IPL and GL in mice ectopically expressing GCaMP6s (Fig. 1k, n-o). Since granule cell dendrites cross the IPL as they project into the EPL, stimulation in the IPL activated GCs and induced Ca^2+^ transients (Fig. 1p). In contrast, after relocation of the stimulation pipette into the GL that is not innervated by GCs, stimulation failed to induce significant Ca^2+^ transients (Fig. 1p-q).

The results show that electrical stimulation in the IPL activates M/T cells and GCs, whereas stimulation in the GL activates M/T cells without stimulation of GCs. Therefore, by relocating the stimulation pipette to different layers it is possible to selectively activate different populations of neurons in the OB.

### Stimulation in the internal plexiform layer induced Ca^2+^ transients in astrocytes in the EPL

To study neurotransmitter-induced changes in astrocytic Ca^2+^ concentration, we used GLAST-Cre^ERT2^ x GCaMP6s^flox^ mice which express GCaMP6s in the majority of astrocytes in the EPL (Fig 2a-d). IPL stimulation of M/T cells and GCs induced Ca^2+^ transients in astrocytes of the EPL (Fig. 2b, e). To ensure astrocytes were responding to neuronal stimulation, only astrocytes at a distance of at least 50 µm from the pipette tip that showed tetrodotoxin (TTX)-sensitive responses were taken into analysis (Fig. s2). We performed a stimulus-response analysis (50 µA, 100 µA, 200 µA and 500 µA) and chose 200 µA, which elicited maximum responses in the astrocytes (n = 26; Fig. 2f-g).

**Figure 2:**
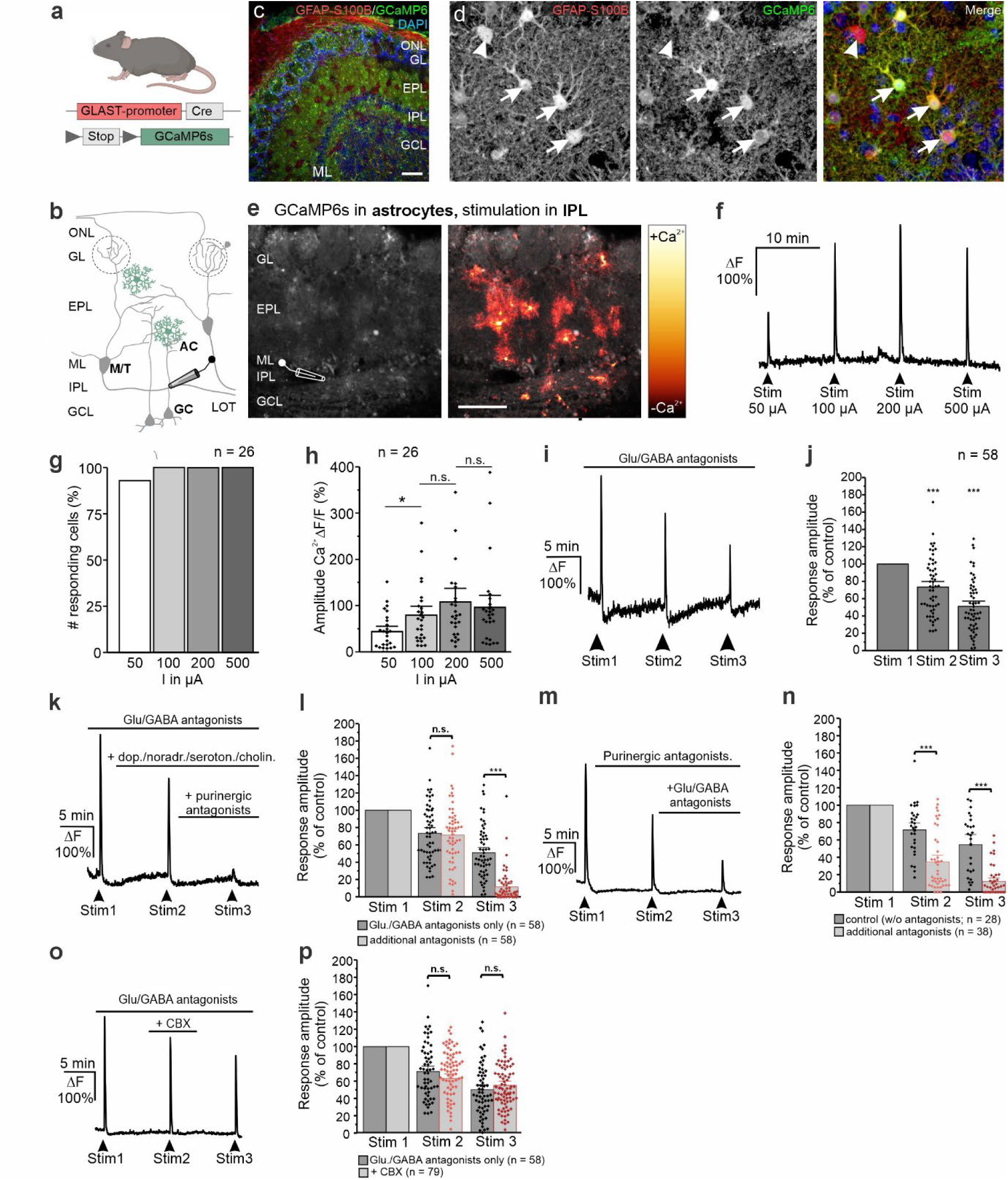
Electrical stimulation of the internal plexiform layer induces purinergic signaling in astrocytes of the external plexiform layer. **(a)** Genetic mouse model expressing GCaMP6s by astrocytes. **(b)** Structure of the OB. GCaMP6s-expressing astrocyte are indicated in green. **(c)** GCaMP6s expression (green) in the OB, astrocytes were immunolabeled against S100b/GFAP (red). Scale: 100 µm. **(d)** Most S100b/GFAP-positive astrocytes expressed GCaMP6s (arrows) with few astrocytes being GCaMP6s-negative (arrowhead). Scale: 10 µm. **(e)** GCaMP6s fluorescence in astrocytes before (left) and after (right) electrical stimulation in the IPL. Scale: 100 µm **(f)** Ca^2+^ transients in astrocytes evoked by stimulation with currents between 50 µA and 500 A. **(g)** Number of responding astrocytes and **(h)** response amplitude. The maximum efficacy was reached with a current amplitude of 200 µA. **(i)** Repetitive stimulation in the IPL leads to a rundown, i.e. Ca^2+^ transients with decreasing amplitude (synaptic isolation by glutamatergic (D-APV, NBQX, MPEP, CPCCOEt) and GABAergic (Gabazine, CGP55845) receptor antagonists). **(j)** Response amplitude upon repetitive stimulation in the IPL in astrocytes normalized to the amplitude of the first stimulation. **(k)** Stimulation-induced Ca^2+^ transients in EPL astrocytes (synaptic isolation, Stim 1). The second stimulation was performed in the additional presence of dopaminergic, noradrenergic, serotonergic and cholinergic antagonists (Stim2; see text for antagonists). Purinergic antagonists (MRS2179, ZM241385) were added during the third stimulation (Stim 3). **(l)** Statistical analysis of the response amplitude (light grey), compared to the respective rundown experiment (dark grey). Antagonizing neuromodulators (Stim2) had no significant effect (n.s.), while purinergic antagonists almost abolished stimulation-induced Ca^2+^ responses (p < 0.001). **(m)** Effect of purinergic antagonists (MRS2179, ZM241385; Stim2) and additional glutamatergic and GABAergic antagonists (NBQX, D-AP5, CPCCOEt, MPEP, gabazine, CGP55845; Stim3) **(n).** Statistical analysis of the response amplitudes in the presence of antagonists (light grey) compared to the rundown experiment (dark grey). Purinergic antagonists (Stim2) significantly reduced the Ca^2+^ transients, and additional glutamatergic/GABAergic antagonists lead to a further reduction (Stim3). ***, p < 0.005. **(o)** Effect of the gap junction hemichannel-inhibitor, carbenoxolone (100 µM CBX; Stim2). (**p)** Statistical analysis of the response amplitude. CBX did not reduce the amplitude of Ca^2+^ transients. n.s., not significant.

To isolate direct neuron-astrocyte transmission and suppress indirect effects mediated by synaptically excited neurons, we inhibited glutamate and GABA receptors in the following experiments. The results show that stimulation in the IPL induced robust Ca^2+^ transients in EPL astrocytes, which were depedent on neuronal activity but were not induced by glutamatergic or GABAergic transmission (206.2 ± 14.4 ΔF, n = 58; Fig 2i-j). As repetitive stimulation induced Ca^2+^ transients in astrocytes with decreasing amplitude, further experiments to investigate which transmitters evoked the astrocytic Ca^2+^ responses were compared with analogous rundown control recordings. The second stimulation-induced Ca^2+^ response in astrocytes was significantly reduced by 26.6 ± 4.3 % of the control (n = 58; p < 0.001), whereas the third stimulation-induced Ca^2+^ response was reduced by 49.0 ± 4.1 % of the control (n = 58; p < 0.001) (Fig. 2i-j). We could exclude bleaching or other technical considerations as the cause of the rundown as Ca^2+^ responses in MCs showed no rundown of Ca^2+^ transients upon repetitive IPL stimulation (Supp. Fig. 1). We previously demonstrated that OB astrocytes respond to neuromodulators such as dopamine, which is released by juxtaglomerular interneurons and norepinephrine, released from terminals arising in the locus coeruleus (Fischer et al., 2021; Fischer et al., 2020). The OB is additionally innervated by neuromodulatory centrifugal fibers, originating from the median raphe nuclei (serotonergic) and the nucleus of the horizontal limb of the diagonal band (cholinergic) (McLean & Shipley, 1987; Rothermel et al., 2014; Shipley et al., 1985). To test whether these neuromodulators contribute to stimulation-induced Ca^2+^ transients in EPL astrocytes, antagonists of adrenergic (α_1_-receptors: prazosine 10 µM; α_2_-receptors: rauwolscine hydrochloride 0.5 µM; β-receptors: ICI118,551 hydrochloride 15 µM), serotonergic (5-HT1 and 5-HT2 receptors: methylsergide maleate 50 µM), cholinergic (mACh-receptors: scopolamine 10 µM; nACh-receptors: mecamylamine 5 µM) and dopaminergic (D1-receptors: SCH23380 20 µM; D2-receptors: sulpiride 50 µM) were applied (Fig. 2k-l). In the presence of these neuromodulatory antagonists, astrocytic Ca^2+^ transients were not reduced compared to Ca^2+^ transients recorded under control conditions (n = 58) (Fig. 2l, Stim 2), suggesting that these neuromodulators do not significantly contribute to Ca^2+^ signals evoked by IPL stimulation in EPL astrocytes.

OB astrocytes respond to ATP and its metabolite adenosine with Ca^2+^ transients via metabotropic P2Y1 receptors (Doengi et al., 2008). Hence, we applied purinergic antagonists (P2Y1 receptors: MRS2179 100 µM; A2A receptors: ZM241385 0.1 µM) in IPL stimulation experiments, which significantly diminished stimulation-induced Ca^2+^ transients in astrocytes to 11.4 ± 2.5 % of the control (p < 0.001) (Fig. 2l, Stim 3). Thus, IPL stimulation releases ATP, which induces Ca^2+^ transients in astrocytes, but not neuromodulators such as dopamine, norephinephrine, serotonin and acetylcholine.

To evaluate whether glutamatergic and GABAergic receptors also mediate stimulation-induced Ca^2+^ transients, we removed the inhibitors and isolated glutamatergic/GABAergic Ca^2+^ signals by inhibiting the P2Y1 and A2A purinergic receptors (Fig. 2m). Inhibition of purinergic receptors now significantly reduced stimulation-induced Ca^2+^ transients in astrocytes by 65.5 ± 5.2 % of the control (n = 38, p < 0.001) (Fig. 2m-n; Stim 2). Additional glutamate and GABA receptor inhibition further reduced the Ca^2+^ transients to 12.4 ± 2.7 % of the control (n = 38, p < 0.001), indicating that astrocytic Ca^2+^ responses to electrical stimulation of M/T cells and GCs are caused by released glutamate and/or GABA (∼ 22%), ATP (∼ 65%) with a small part of the response unaccounted for (∼ 12 %) (Fig. 2m-n, Stim 3).

Our results suggest that release of ATP from M/T cell and/or granule cell dendrites account for the largest part of Ca^2+^ transients in astrocytes in the EPL. Another source of ATP in the OB are astrocytes themselves, releasing ATP through gap junction hemichannels (Roux et al., 2015). Hence, ATP-dependent activation of astrocytes might trigger release of additional ATP, boosting Ca^2+^ transients in neighboring astrocytes. This boosting proved not to be the case as carbenoxolone (CBX, 100 µM), a gap junction and hemichannel inhibitor, did not reduce the stimulation-induced Ca^2+^ transients in astrocytes (Fig. 2o-p).

### Activation of granule cells initiates ATP release and Ca^2+^ transients in astrocytes in the EPL

Within the EPL axonal projections are largely lacking and most synapses are dendrodendritic synapses between dendrites of M/T cells and GCs, which are the likely source of ATP. To identify which cells release the ATP we compared the astrocytic Ca^2+^ transients in response to stimulation of both M/T cells and GCs (IPL stimulation) or M/T cells alone (GL stimulation, see Fig. 1; Fig. 3a-b). Electrical stimulation in the IPL induced Ca^2+^ transients in 89% of all astrocytes in the field of view (Fig. 3 c, e-f). In contrast, stimulation in the GL evoked Ca^2+^ transients in only 22% of the astrocytes, half of which also responded to IPL stimulation (11%) with the remaining 11% only responding to stimulation in the GL (Fig. 3d, e-f). Not only were many fewer astrocytes responding to GL stimulation but the Ca^2+^ transients were also significantly smaller than those evoked by IPL stimulation (13.9 ± 4.8 % ΔF; n = 21 vs 74.6 ± 21.0 % ΔF; n = 18; p < 0.001, Fig. 3g). Thus M/T cells alone do not strongly induce Ca^2+^ responses in EPL astrocytes, whereas the activation of interneurons, in particular GCs, by IPL stimulation drives most of the astrocytic Ca^2+^ responses. To visualize ATP release from GCs, we expressed the genetically encoded P2Y1 receptor-based sensor for ATP, GRAB-ATP1.0 in astrocytes (Wu et al., 2022)(Fig. 3h-j). We calibrated the GRAB-ATP1.0 signal by applying increasing concentrations of ATP from 3-100 µM via the perfusion system and found a clear dose-response relationship, rising from 163.57 ± 37.0 % ΔF at 3 µM ATP to 409.12 ± 59.51 % ΔF at 100 µM ATP (Fig. 3k, l). Electrical stimulation in the IPL induced a robust increase in extracellular ATP within the EPL with an amplitude of 138.11 ± 16.2 % ΔF, whereas stimulation in the GL elicited significantly smaller ATP signals (4.4 ± 2.5 % ΔF, n = 28; p < 0.001; Fig. 3k, m). The results demonstrate that activation of GCs leads to ATP release in the EPL, whereas activation of M/T cells alone is insufficient to trigger ATP release in the EPL.

**Figure 3:**
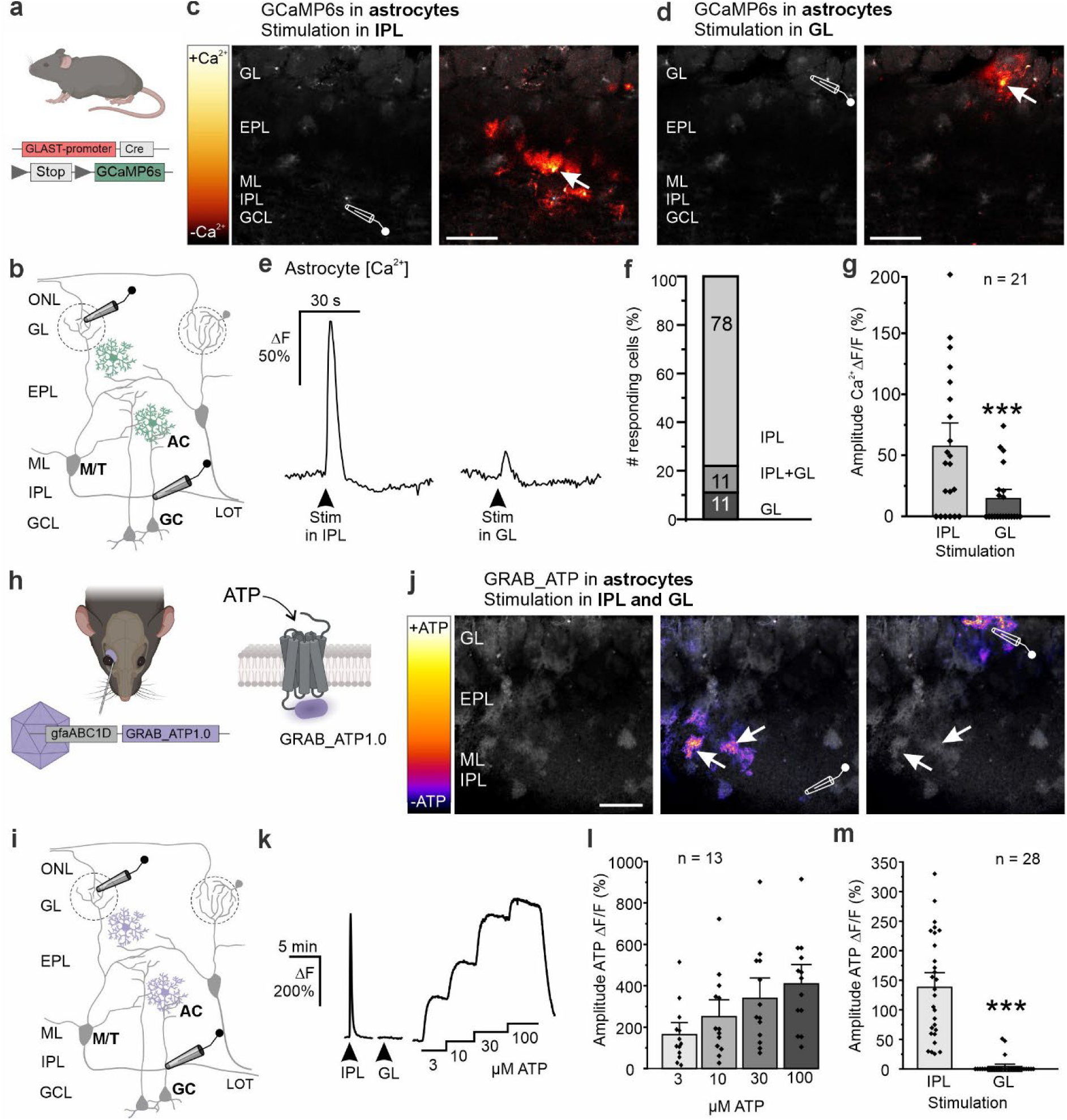
Stimulation of granule cells induces ATP release and Ca^2+^ transients in astrocytes of the external plexiform layer. **(a)** Genetic mouse model expressing GcaMP6s in astrocytes. **(b)** OB with GCaMP6s-expressing astrocyte indicated in green. **(c)** Image sequence of GCaMP6s fluorescence in astrocytes before (left) and after (right) electrical stimulation in the IPL. Scale: 100 µm. Stimulation in the IPL resulted in Ca^2+^ transients in numerous astrocytes in the EPL (arrow), while stimulation in the GL of the same preparation **(d)** failed to trigger Ca^2+^ responses in astrocytes except for those in the direct vicinity of the stimulation pipette (arrow). **(e)** Ca^2+^ transients induced by stimulation in IPL (left) compared to stimulation in GL (right). **(f)** Proportion of responding astrocytes to different stimulation positions. The majority of astrocytes responded to IPL stimulation and IPL as well as GL stimulation, while few astrocytes responded only to GL stimulation. **(g)** Ca^2+^ transients upon stimulation in the IPL were significantly larger than upon stimulation in the GL. **(h)** AAVs carrying GRAB-ATP1.0 were injected into the retro-bulbar sinus, leading to expression of the GRAB-ATP1.0 sensor in astrocytes to visualize extracellular ATP. **(i)** OB with GRAB-ATP1.0-expressing astrocytes indicated in violet. **(j)** Image sequence of GRAB-ATP1.0 fluorescence before (left) and after (right) electrical stimulation in the IPL, resulting in ATP release in the EPL (arrows), and GL. Scale: 100 µm. **(k)** Representative ATP-signal in response to IPL and GL stimulation. Calibration of the GRAB-ATP1.0 sensor using ascending concentrations of ATP applied via the perfusion system (3, 10, 30, 100 µM ATP). **(l)** Calibration of the GRAB-ATP1.0 sensor expressed in astrocytes. ATP signals in astrocytes increase with ascending ATP concentrations applied via the perfusion system. ***, p < 0.001. Scale: 100 µm. **(m)** Response amplitude of stimulation-induced ATP signals in astrocytes.

### Optogenetic stimulation but not synaptic stimulation of granule cells excites astrocytes

As mentioned, it is not possible to electrically stimulate only GCs. We therefore used mice expressing channelrhodopsin-2 (ChR2) to drive action potential firing in either GCs or M/T cells (Fig. 4a-b; 4h-i). In Pcdh21-Cre x ChR2^flox^ mice, ChR2 was expressed exclusively in M/T cells (entopic expression) in five out of ten mice (Fig. 4h-i; Fig.s3). In their littermates ChR2 was expressed in GCs and other inhibitory interneurons (ectopic expression) (Fig. s3). Stimulation was achieved by applying repetitive light flashes (5 ms, 50 Hz, train of 7.5-10 s) using a blue LED (peak wavelength: 450 nm), which elicited a single action potential per flash and hence bursts of actions potentials at 50 Hz in ChR2-expressing GCs and M/T cells, respectively (Fig. 4 c, j). The red fluorescent Ca^2+^ indicator Rhod-2 was employed to image Ca^2+^ signals in astrocytes (Fig. 4b, i). Since in the OB only astrocytes but not neurons respond to bath application of ADP with Ca^2+^ increases (Doengi et al., 2008; Fischer et al., 2020), astrocytes were identified by application of ADP. In our experiments, 85% of Rhod-2-loaded cells responded to ADP and hence were identified as astrocytes (Fig. 4d, k).

**Figure 4:**
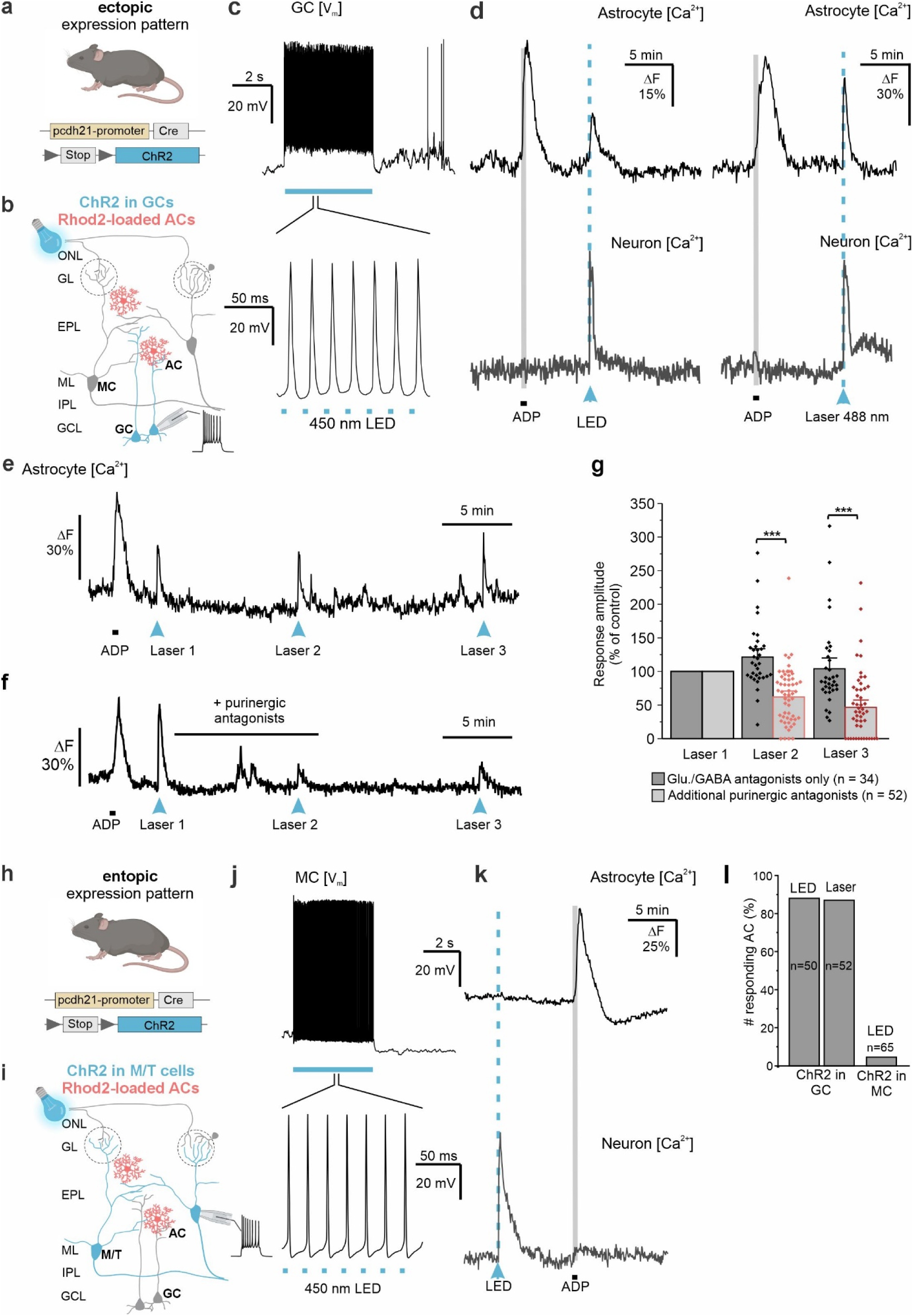
Photostimulation of ChR2-expressing granule cells activates astrocytes in the EPL. **(a)** Mouse model ectopically expressing ChR2 controlled by the pcdh21-promoter. **(b)** OB with ChR2-expressing GCs indicated in blue. Astrocytes and neurons were loaded with Rhod2-AM (indicated in red for astrocytes). ChR2-expressing GCs were excited by illumination with blue light (LED; 5-ms light pulses, 50 Hz, 10 s). **(c)** Recordings of GC membrane potential (Vm) upon blue light illumination. Photostimulation of ChR2 in GCs leads to depolarization and action potential firing in GCs, as well as Ca^2+^ responses in astrocytes and neurons in the EPL **(d)**. ADP was bath-applied to identify astrocytes (synaptic isolation: glutamatergic D-APV, NBQX, MPEP, CPCCOEt and GABAergic antagonists gabazine, CGP55845). **(d)** GCs expressing ChR2 were stimulated using a laser of 488 nm wavelength that was directed exclusively to the GCL and Ca^2+^ responses were recorded in astrocytes and neurons. **(e)** Repetitive photostimulation of GCs (in the presence of glutamatergic and GABAergic antagonists) evoked Ca^2+^ transients of constant amplitude. **(f)** Additional antagonists of purinergic receptors (100 µM MRS2179; 0.1 µM ZM241385) attenuated Ca^2+^ responses in astrocytes evoked by optogenetic stimulation of GCs. **(g)** Analysis of the effect of purinergic blockage on astrocytic Ca^2+^ transients evoked by repetitive optogenetic stimulation of GCs. ***, p < 0.001. **(h)** Mouse model expressing ChR2 controlled by the pcdh21-promoter. **(i)** OB with ChR2-expressing M/T cells indicated in blue. M/T cells were excited by illumination with blue light (LED; 5-ms light pulses, 50 Hz, 10 s). **(j)** Recordings of MC membrane potential (Vm) and action potentials upon blue light illumination. **(k)** Optogenetic stimulation of M/T cells failed to induce Ca^2+^ signals in astrocytes. **(l)** Fraction of astrocytes responding to optogenetic stimulation of GCs (by LED and laser) as well as M/T cells.

Wide-field optogenetic stimulation of ChR2-expressing interneurons, including GCs, in the presence of glutamatergic and GABAergic antagonists induced Ca^2+^ transients in astrocytes in the EPL (n = 50) (Fig. 4d). To excite GCs more specifically, we directed the 488-nm laser of a confocal microscope to the granule cell layer and scanned a square of approximately 300 x 100 µm, sparing other layers of the OB and hence avoiding stimulation of interneurons in the glomerular and external plexiform layers, which might also express ChR2. Stimulation of ChR2-expressing GCs with the scanning laser for a period of 10 seconds evoked a Ca^2+^ increase in astrocytes in the EPL (n = 53) (Fig. 4d). Repeated optogenetic stimulation of GCs (10 min interval) produced Ca^2+^ transients in Rhod-2-loaded astrocytes in the EPL with similar amplitude (Fig. 4e). Purinergic antagonists (MRS2179 100 µM; ZM241385 0.1 µM) significantly reduced the ampitude of Ca^2+^ transients evoked by laser illumination of the granule cell layer by 41.4 ± 4.8 % (n = 51), confirming prominent contribution of purinergic signaling in GC-astrocyte communication (Fig. 4f-g; Laser 2). The effect of purinergic antagonists was irreversible after wash out for 10 minutes (Fig. 4f-g; laser 3). Finally, we were interested whether not only optogenetic stimulation of GCs, but also synaptic excitation of GCs by glutamate released from M/T cells at dendrodendritic synapses in the EPL leads to ATP release from GCs and astrocyte activation. To evoke synaptic granule cell excitation, we optogenetically stimulated ChR2-expressing M/T cells in the absence of any neurotransmitter receptor antagonist and visualized changes in Ca^2+^ in astrocytes using Rhod-2 (Fig. 4h, k). Surprisingly, when GCs were synaptically excited by M/T cells, they failed to trigger Ca^2+^ transients in EPL astrocytes (n = 65), whereas interneurons in the GL and/or EPL responded with Ca^2+^ signals (n = 11) (Fig. 4k), confirming glutamate release from M/T cells. In summary, an average of 88 % and 87 % of astrocytes responded to optogenetic stimulation of GCs with LED and laser, respectively, whereas only three out of 65 astrocytes (5 %) responded to synaptic excitation of GCs evoked by optogenetic M/T cell stimulation (Fig. 4l).

### Top-down glutamatergic input from aPC neurons drives granule cell firing resulting in ATP-dependent Ca^2+^ signaling in astrocytes

To address the question why GCs excite astrocytes when optogenetically stimulated but fail to excite astrocytes when synaptically stimulated by M/T cells, we recorded GC membrane potentials using whole-cell patch-clamp recordings while stimulating ChR2 expressing M/T cells (5 ms blue LED light pulses, 50 Hz, 10 s) (Fig. 5a). During M/T cell stimulation, GCs (8 cells from 3 animals) received a burst of excitatory postsynaptic potentials that added up to a steady state depolarization as long as optogenetic stimulation of M/T cells lasted (Fig. 5b). However, only one of the recorded GCs fired a long-lasting burst of action potentials (Fig. 5, GC type 3), whereas no action potentials were evoked in most of the GCs (5 out of 8) (Fig. 5b, GC type 1). In the remaining 2 cells, one action potential was recorded (Fig. 5b, GC type 2). This firing behavior was in marked contrast to optogenetically stimulated GCs that maintained action potential firing for the entire duration of stimulation (Fig. 4c). Thus, synaptic input from M/T cellsis not sufficient to drive dendritic ATP release from GCs. However, GCs not only receive synaptic input from M/T cells, but also from glutamatergic pyramidal cells of the aPC. This cortical input regulates synaptic integration in GCs, resulting in strong excitatory postsynaptic potentials (EPSPs) (Balu et al., 2007; Pressler & Strowbridge, 2017). To validate, whether electrical stimulation of glutamatergic fibers, projecting from the aPC results in long-lasting depolarization and firing in GCs, comparable to optogenetic stimulation (Fig. 4c), we adapted a previously published stimulation protocol to electrical stimulate aPC input into the OB. Therefore, we placed a stimulation pipette in the deeper part of the granule cell layer to stimulate aPC fibers in a distance of approximately 50-100 µm to the proximal dendrite of the recorded GC and recorded its membrane potential (Fig. 5c-d) (Balu et al., 2007). In all recorded cells (n = 8), electrical stimulation of cortical fibers induced a long-lasting burst of action potentials that lasted throughout the stimulation period (Fig. 5d). Next, we investigated whether electrical stimulation of cortical fiber projections is sufficient to drive ATP release of GCs, inducing Ca^2+^ signaling in EPL astrocytes. To avoid direct stimulation of GCs, we placed the stimulation pipette in the aPC, using an in-toto preparation containing both the OB and the aPC (Fig. 5e). Top-down projections from the aPC to the OB are sparse and dissuse, thus astrocytes that were within the reach of GCs excited by aPC stimulation were found in very few cases only (n = 3 out of 12 trials/8 animals). Electrical stimulation of cortical fiber projections in the aPC induced Ca^2+^ transients in astrocytes of the EPL, indicating that glutamatergic cortical input into the OB drives granule cell firing, subsequent ATP release and astrocyte activity (Fig. 5f-g). In summary, Ca^2+^ transients in EPL astrocytes are driven by ATP released from GCs that greatly depend on the origin of GC synaptic activation.

**Figure 5:**
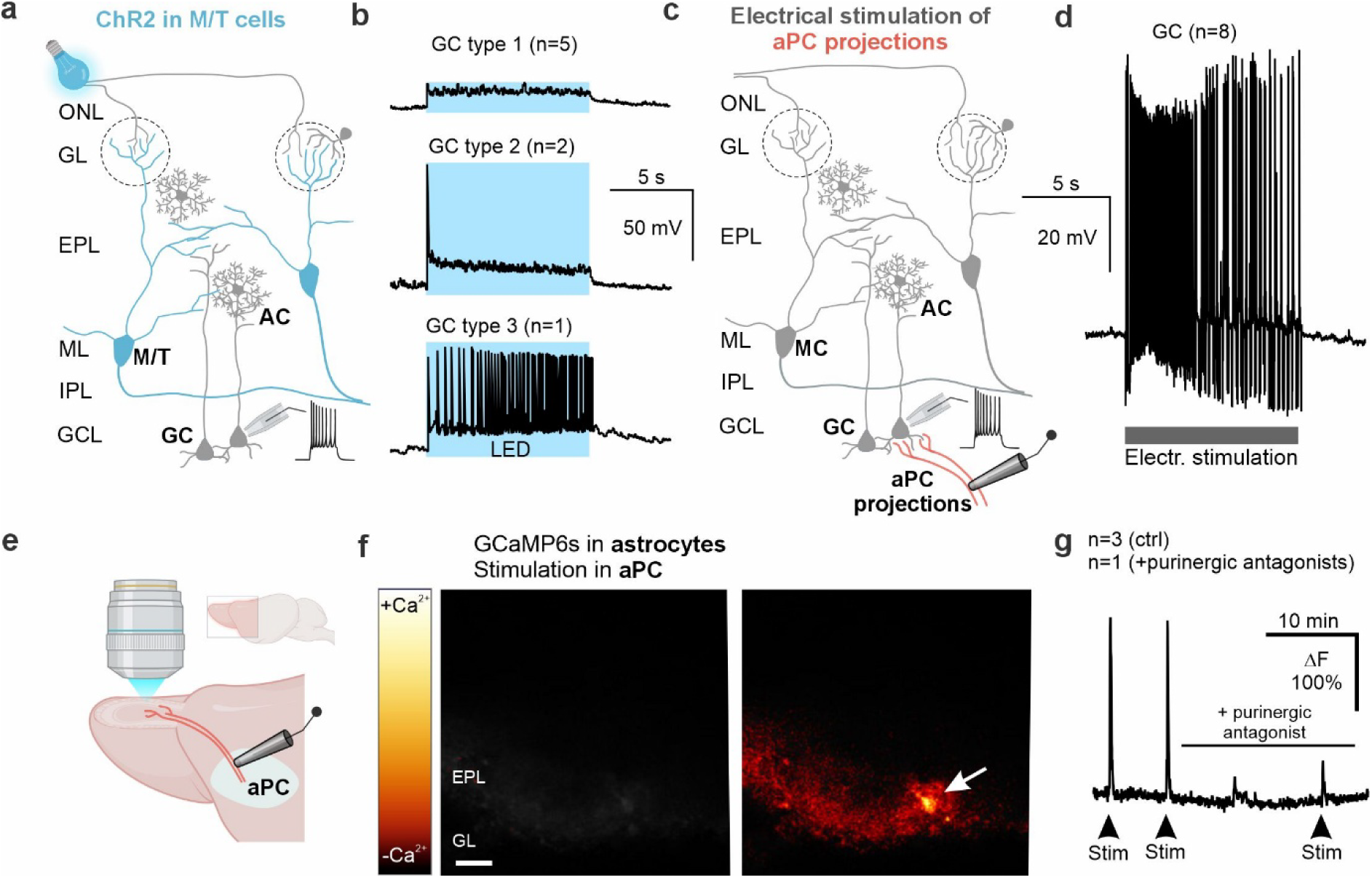
Cortical top-down projections drive granule cell firing and P2Y1-mediated Ca^2+^ signaling in astrocytes. **(a)** Illustration of the experimental paradigm. Optogenetic stimulation of M/T cells (blue) results in distinct depolarization patterns in GCs, as assessed by patch-clamp recordings **(b)** Single-cell electrophysiological recordings of GCs, responding to optogenetic M/T cell stimulation (LED; 5-ms light pulses, 50 Hz, 10 s) with depolarization, either without (GC type 1; 5 out of 8 cells) or with firing of single action potentials (GC type 2; 2 out of 8 cells). Only in one recorded cell, a long-lasting burst of action potentials was elicited (GC type 3; 1 out of 8 cells). **(c)** Illustration of electrical stimulation of projection fibers from the anterior piriform cortex (aPC) in wildtype mice and simultaneous recordings of granule cell membrane potential. **(d)** Electrical stimulation of aPC projections resulted in long-lasting action potential firing of GCs lasting throughout the stimulation period (n = 8). **(e)** Electrical stimulation of aPC projection fibers in a in-toto preparation containing the OB and the aPC and Ca^2+^ imaging in GCaMP6s-expressing astrocytes of the EPL. **(f)** Image sequence of Ca^2+^ transients in astrocytes before (left) and after (right) electrical stimulation of the aPC (arrow). Scale: 100 µm. **(g)** Example trace of stimulation-induced Ca^2+^ transients in astrocytes, showing repetitiv activation (Stim 1 and Stim 2) and a great reduction in the presence of the P2Y1 receptor antagonist (Stim 3; MRS2179, 100 µM). n = 3 for control conditions; n = 1 for purinergic inhibition.

## Discussion

How cortical top-down modulation contributes to astrocyte activation in lower brain areas and whether astrocytes are able to distinguish between bottom-up and top-down pathways has not been investigated so far. The aim of our study was to unravel how astrocytes are embedded in the neuronal network in the OB EPL that is highly regulated by both bottom-up and top-down information processing. Due to the specific cellular organisation of the OB, we were able to selectively stimulate different neuron populations and investigate the respective astrocyte responses, addressing the question whether astrocytes respond to neuronal activity in a general or in a cell type-specific manner. Our results show that EPL astrocytes respond to ATP released by GC firing elicited by aPC top-down projections. M/T cells, in contrast, appear not to excite astrocytes by release of, for example, ATP or glutamate.

### Neuronal excitation of astrocytes in the EPL is cell type-specific

We used cell type-specific transgenic mouse models, different stimulation protocols, cell-type specific optogenetics and pharmacological tools to dissect the components of the tripartite synapse in the EPL of the OB. We established stimulation protocols employing a Pcdh21-Cre mouse line that is supposed to drive Cre expression in OB principal neurons, but not interneurons. In some of the offspring, however, Cre-dependent expression of the Ca^2+^ sensor GCaMP6s or of ChR2 was detected exclusively in interneurons such as GCs. We made this observation using the Pcdh21-Cre driver line in the present study (Tg(Cdhr1-cre)KG66Gsat/Mmucd; Gensat) as well as an independently developed Pcdh21-Cre mouse line (C57BL/6Cr-Tg(Pcdh21-cre)BYoko; Riken; data not shown) (Nagai et al., 2005). The entopic and ectopic Cre-dependent expression of GCaMP6s enabled us to assess the efficiency of electrical stimulation in both, M/T cells and GCs. This allowed for the development of stimulation protocols distinguishing between M/T cell-to-astrocyte and GC-to-astrocyte transmission. In addition, we separately photostimulated M/T cells and GCs expressing ChR2. Both electrical and optogenetic stimulation of M/T cells failed to evoke considerable Ca^2+^ signals in astrocytes, while stimulation of GCs triggered large Ca^2+^ transients in the majority of astrocytes in the EPL. Astrocytes in virtually all brain regions express P2Y receptors (Verkhratsky & Nedergaard, 2018), while the situation in the OB is more complex, since here astrocytes additionally express A2A adenosine receptors (Doengi et al., 2008). Thus, enzymatic breakdown of ATP released from GCs does not result in inactivation of the transmission process, as for other neurotransmitters, but results in activation of the even more sensitive adenosinergic signaling. Therefore, P2Y1 and A2A receptors were blocked to test purinergic transmission between GCs and astrocytes. In addition to purinergic transmission, we tested the contribution of glutamatergic and GABAergic transmission. Unexpectedly, glutamate release from M/T cells failed to activate Ca^2+^ signaling in astrocytes upon optogenetic stimulation of M/T cells, which was done in the absence of glutamatergic antagonists. OB astrocytes express mGluR_5_ in juvenile mice, however, mGluR_5_ disappears in OB astrocytes after the first 2-3 weeks after birth (Beiersdorfer et al., 2019; Otsu et al., 2015) as shown in cortical astrocytes before (Sun et al., 2013). We investigated astrocytes in mice at an age of approximately 5 weeks, which hence lack mGluR_5_, but have been shown to generate Ca^2+^ signals by recruiting Ca^2+^-permeable AMPA receptors (Droste et al., 2017). Our results suggest that Ca^2+^-permeable AMPA receptors in EPL astrocytes are not activated by glutamate released from M/T cells, however, we cannot rule out astrocyte activation by glutamate released from aPC top-down projections. Another striking result is that stimulation of M/T cells also failed to activate astrocytes indirectly by excitation of GCs. GCs are excited by glutamate released from M/T cells at reciprocal dendrodendritic synapses, which leads to release of GABA from GCs, but apparently fails to trigger ATP release that would activate astrocytes. The discrepancy between direct stimulation of GCs evoking Ca^2+^ transients in astrocytes and M/T cell-dependent synaptic stimulation of GCs failing to evoke Ca^2+^ transients in astrocytes can be explained by the specific anatomical and physiological properties of the granule cell’s dendritic spines. As part of the reciprocal synapse between M/T cells and GCs, the spine consists of a large spine head and a thin elongated spine neck with high electrical resistance, which enables the spine to act independently of the remaining GC dendrite (Egger & Kuner, 2021; Lage-Rupprecht et al., 2020). Hence, excitation of GC spines by M/T cells might not be sufficient to excite dendritic compartments that comprise ATP release sites, while triggering action potentials, as accomplished by electrical or optogenetic stimulation of GCs, covers the entire granule cell dendrite including ATP release sites. Indeed, optogenetic stimulation of a large number of M/T cells failed to elicit multiple action potentials in GCs, supporting the hypothesis that reciprocal dendrodendritic synapses build microcircuits that act independently but do not add up sufficiently to trigger substantial action potential firing in GCs, essential to evoke ATP release. This provides a hitherto undescribed complexity of neuron-astrocyte interaction that is not only cell type-specific but also dependent on the firing behavior of GCs. While dendrodendritic excitation of GCs by M/T cells failed to evoke a large number of action potentials, excitation of GCs by glutamatergic synaptic inputs by top-down innervations originating from pyramidal cells in the aPC could elicit trains of action potentials, in particular when occuring simultaneously with M/T cell excitation (Balu et al., 2007)(Fig. 6).

**Figure 6:**
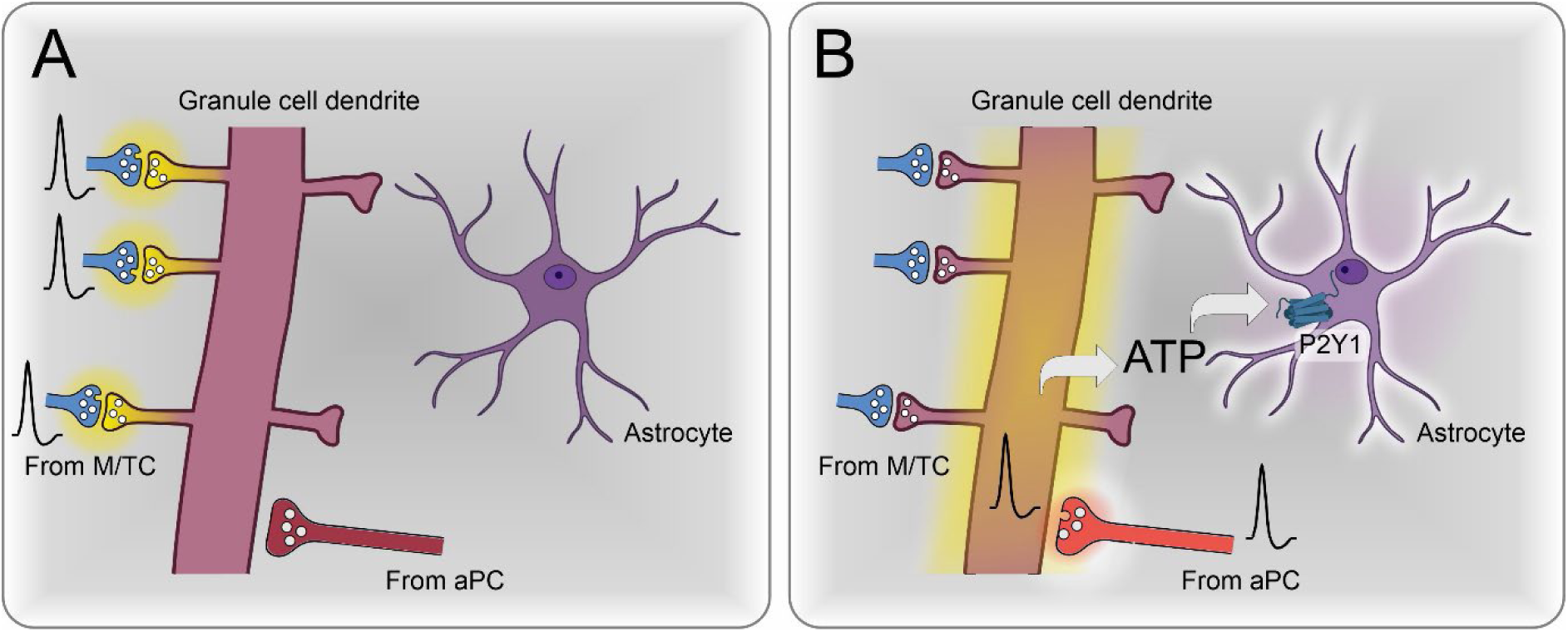
Schematic model of granule cell to astrocyte interaction in the OB. **(a)** Excitation of dendrodendritic synapses at granule cell dendritic spines by M/T cells leads to local depolarization of spines. **(b)** Excitation of GCs by feedback projections from pyramidal cells in the anterior piriform cortex (aPC) leads to global depolarization of the granule cell that is sufficient to trigger action potentials and ATP release. ATP activates astrocytic P2Y1 receptors resulting in Ca^2+^ transients.

### Astrocytes in information processing in the olfactory bulb

The piriform cortex is involved in odor identification both in rodents and in humans, but also functions as an associative cortex (Beiersdorfer et al., 2024; Gottfried, 2010; Kehl et al., 2024; Pashkovski et al., 2020). The aPC receives input from the OB by M/T cell projections and hence is activated during odor perception. However, the aPC is most robustly active when olfactory information needs to be transformed into behaviorally relevant, often associative, representations—especially under conditions of odor-driven learning, stress/survival significance, internal nutrient state changes, and social cue processing. It becomes strongly active when odors must be associated with meaning, value, or context, rather than merely detected (Roth & Sullivan, 2005). The aPC shows increased immediate early gene expression (e.g., c-fos) when animals are exposed to predator odors that induce robust innate fear and stress responses. Under these conditions, aPC activity is coupled to beta-band oscillatory input from the OB and participates in a wider network (bed nucleus of the stria terminalis, amygdalopiriform area, HPA axis) that links odor cues to endocrine and behavioral stress outputs (Matsukawa et al., 2022). Recent work shows that aPC neurons exhibit distinct firing patterns and calcium transients in response to familiar versus novel conspecifics, indicating that social odor cues engage the aPC in a familiarity- and novelty-dependent manner. This social context–dependent activation is essential for normal social recognition and preference behaviors, positioning the aPC as a hub where olfactory and social information converge (Zhou et al., 2025). In addition, the aPC is modulated by learning, anticipation and contextual cues (Kadohisa & Wilson, 2006; Mediavilla et al., 2016; Roesch et al., 2007). The examples stated above demonstrate that aPC activity and its top-down modulation of the OB are strongly regulated by context and brain state, thus our results predict a context- and state-dependent contribution of astrocytes in the EPL to odor information processing. With their fine and densely branched processes, astrocytes cover almost the entire parenchyma of the OB (Klein et al., 2020). Knock out of the transcription factor SOX9 in OB astrocytes results in severe morphological and physiological alterations of the astrocytes, accompanied by impairment of M/T cells and behavioral deficits such as changes in odor detection sensitivity and odor discrimination (Ung et al., 2021). Astrocytes release glutamate and GABA as gliotransmitters (Kozlov et al., 2006). In addition, astrocytes in the GL release ATP that, after degradation to adenosine, affects the firing pattern of MCs and, consequently, the output information of the OB propagated to the olfactory cortex (Rotermund et al., 2018; Roux et al., 2015). Therefore, GC-derived ATP and the subsequent Ca²⁺ signaling in EPL astrocytes may also modulate neuronal output through gliotransmitter release at the lateral dendrites of principal neurons in the EPL. However, OB astrocytes not only transmit information to neurons, but also to other types of glial cells. For instance, astrocytes and olfactory ensheathing cells are coupled by gap junctions in panglial networks that propagate Ca^2+^ waves from the glomerular layer to the nerve layer and vice versa (Beiersdorfer et al., 2019). In addition, astrocytes mediate neurovascular coupling, thus adjusting vessel diameter and rate of blood flow (Beiersdorfer et al., 2020; Doengi et al., 2009; Otsu et al., 2015; Petzold et al., 2008). In conclusion, astrocytes in the OB are critical for neuronal performance and homeostasis, but how the intricate interaction between GCs and astrocytes in the EPL contributes to these functions is still enigmatic and needs to be addressed in future studies.

## Materials and Methods

### Animals and olfactory bulb preparation

Colonies of GLAST-Cre^ERT2^ (Slc1a3^tm1(cre/ERT2)Mgoe^), Pcdh21-Cre (Tg(Cdhr1-cre)KG66Gsat/Mmucd; Gensat, Rockefeller University, New York), GCaMP6s^flox^ (Ai96, The Jackson Laboratory, Bar Harbor, ME USA) and ChR2^flox^ (B6;129S-Gt(ROSA)26Sor^tm32(CAG-COP4*H134R/EYFP)Hze^/J; The Jackson Laboratory, Bar Habor, Maine) were obtained from the institutional animal facility at the University of Hamburg. Pcdh21-Cre founder mice were generously provided by Markus Rothermel (Aachen, Germany), GLAST-Cre^ERT2^ by Frank Kirchhoff (Homburg, Germany), ChR2^flox^ by Franziska Schneider-Warme (Freiburg, Germany). Animal rearing and all experimental procedures were performed according to the European Union’s guidelines and approved by local animal welfare authorities (N 107/2019; Behörde für Justiz und Verbraucherschutz, Hamburg, Germany). To induce GCaMP6s expression by Cre in GLAST-Cre^ERT2^ x GCaMP6s^flox^ mice, tamoxifen (Carbolution, St Ingbert, Germany) was dissolved in ethanol and miglyol and injected intraperitoneally for three consecutive days (starting p21; 100 mg/kg bodyweight). Animals were analyzed seven to twelve days after the first injection. OB horizontal slices were prepared as described before (Schulz et al. 2018) and stored for 30 min in carbogen-gassed ACSF at 30°C for recovery. Preparation of the OB in-totos was modified from previous descriptions (Stavermann et al., 2012). In brief, both OBs with the rostral part of the cortex attached (including the piriform cortex) were removed from the opened head and glued to the stage of a vibratome (VT1200, Leica, Germany), the lateral view of the bulb facing upwards. A slice of approximately 300 µm was cut from the top of the OB to expose the surface of a sagittal cross section including all layers, while the cortex and projections from the cortex to the OB were left intact. In-toto preparations were stored for 30 min in carbogen-gassed ACSF at 30°C for recovery. For experiments, in-toto preparations were glued onto a round coverslip (10 mm diameter), the surface of the sagittal section facing upwards, and transferred to an experimental chamber. Standard artificial cerebrospinal fluid (ACSF) consisted of (in mM): 120 NaCl, 2.5 KCl, 1 NaH_2_PO_4_x2H_2_O, 26 NaHCO_3_, 2.8 D-(+)-glucose, 1 MgCl_2_, 2 CaCl_2_. The preparation solution consisted of (in mM) 83 NaCl, 1 NaH_2_PO_4_x2H_2_O, 26.2 NaHCO_3_, 2.5 KCl, 70 sucrose, 20 D-(+)-glucose, 2.5 MgSO_4_ x7 H_2_O. Both solutions were continuously gassed with carbogen (95% O_2_, 5% CO_2_) to maintain the pH of 7.4 and to ensure oxygen supply.

### Virus injection of GRAB-ATP1.0

To express the extracellular ATP sensor (GRAB-ATP1.0) specifically in astrocytes, we used the plasmid: pAAV-gfaABC1D-GRAB_ATP1.0 (Addgene #167579), generously provided as a gift from Yulong Li which was packaged into AAV2/PhP.AX (pUCmini-iCAP-PHP.AX, Addgene #195218), was generously provided as a gift by Viviana Gradinaru (Jang et al., 2023) at the UKE Vector Facility. Virus suspensions were diluted with sterile saline and injected i.v. into the retro-bulbar sinus of anesthetized mice. Each animal received an injection of 70 µl virus suspension, containing 1 x 10^11^ viral genomes. Animals were analyzed three weeks after i.v. injection.

### Immunohistochemistry

After dissection of OBs, 220 µm thick horizontal slices were prepared using a vibratome (Leica VT 1000 S, Leica Biosystems, Wetzlar, Germany). The slices were kept in formalin (4% paraformaldehyde in PBS, pH 7.4) for 1 h at room temperature (RT), followed by the incubation in blocking solution (10% normal goat serum (NGS), 0.5% Triton X-100 in PBS) for 1 h at RT. Afterwards, the slices were incubated for 48 h at 4°C with the following primary antibodies: rabbit anti-S100B (Dako, Hamburg, Germany; 1:1000); rabbit anti-GFAP (Dako, 1:1000); chicken anti-GFP (which also labels GCaMP6s and YFP; Novus; 1:500). The antibodies were diluted in 1% NGS, 0.05% TritonX100 in PBS. Slices were incubated in PBS with the following secondary antibodies for 24 h at 4°C: goat anti-rabbit Alexa 555 (Invitrogen; 1:1000); goat anti-chicken Alexa 488 (Abcam; 1:1000). Additionally, Dapi (5 µM; BioChemica) or Hoechst 33342 (10 µM; Molecular Probes) was added to stain nuclei. The slices were mounted on slides using self-hardening embedding medium (Immu-Mount, Thermo Fisher). Immunohistological stainings were analyzed using a confocal microscope (Nikon eC1). Confocal images were adjusted for contrast and brightness using Adobe Photoshop CS6 and ImageJ.

### Ca^2+^ imaging and data analysis

OB slices were transferred into a recording chamber and secured with a U-shaped platinum wire with nylon strings. Changes of the cytosolic Ca^2+^ concentration in either astrocytes (GLAST-Cre^ERT2^-driven GCaMP6s expression)(Lohr et al., 2021) or neurons (Pcdh21-Cre-driven GCaMP6s expression) was detected by changes in fluorescence of GCaMP6s (excitation: 488 nm) using a confocal microscope (eC1, Nikon, Düsseldorf, Germany). Images were acquired at a time rate of one frame every 1.5 s. All drugs were applied via the perfusion system with an incubation period of at least 10 min prior to electrical stimulation. For electrical stimulation of axons and dendrites a glass pipette with a resistance of 1-2.5 MΩ filled with ACSF was placed in the IPL or GL. The stimulation protocol consisted of 10 cycles, with 20 single pulses each cycle and a single pulse duration of 100 µs at 100 Hz frequency. One cycle was followed by a 300 ms non-stimulation phase. 50-500 µA stimulation currents were applied. The total length of the stimulation protocol was 5 s. This stimulation protocol is also referred to as “sniff cycle stimulation”, as it simulates rhythmic inhalation of odors and the corresponding neuronal activity within the OB (Kepecs, Uchida et al. 2006). To analyze changes in single cell somata or processes, regions of interest were defined using Nikon EZ-C1 3.90 software. Cells chosen for analysis had a minimum distance radius of 50 μm from the stimulation pipette to exclude cells that might be directly activated by the stimulation current. In addition, 1 µM tetrodotoxin (TTX) was applied at the end of each experiment to suppress action potential firing and astrocytes that responded in the presence of TTX were considered as directly stimulated and were discarded (Fig. s2). In experiments using optogenetic stimulation, Ca^2+^ was recorded using Rhod-2 as Ca^2+^ indicator. Slices were incubated in 5 µM Rhod-2 in ACSF for 30-45 minutes. For electrical stimulation of aPC neurons, in-toto preparations containing the cortex and OB from GLAST-Cre^ERT2^ x GCaMP6s^fl^ mice were used. A glass pipette with a resistance of 0.8-1 MΩ, filled with ACSF, was positioned in the anterior piriform cortex to deliver electrical stimuli according to the “sniff cycle” protocol. Optogenetic stimulation was achieved by applying repetitive light flashes (5 ms, 50 Hz, train of 7.5-10 s) using a blue LED (peak wavelength: 450 nm) or by using the 488-nm scanning laser of the confocal microscope. Changes in cytosolic Ca^2+^ were recorded as relative changes in GCaMP6s or Rhod-2 fluorescence (ΔF) with respect to the resting fluorescence, which was normalized to 100%. Quantification of the Ca^2+^ transients was established by the amplitude calculation of ΔF. All values are stated as mean values ± standard error of the mean. The number of experiments is given as n = x, whereas x is the number of analyzed cells. For every set of experiment more than 3 animals were analyzed. Statistical significance was assessed by comparing the mean of the control experiment (rundown experiment) to the mean of the actual experiment using the Mann-Whitney U-Test with the error probability p (* p < 0.05; ** p < 0.01; *** p < 0.001).

### Electrophysiological Recordings

Mitral and granule cells of the main OB were investigated using the patch-clamp technique (Multi Clamp 700 B amplifier and pCLAMP 10 software, Molecular Devices, Biberach, Germany, and EPC10 with Patchmaster software, HEKA, Lambrecht, Germany). During the experiments, brain slices were continuously superfused with ACSF. Whole-cell recordings of mitral cells were performed using patch pipettes with a resistance of ∼3 MΩ, while pipettes with a resistance of ∼5-6 MΩ were used for granule cells. The pipette solution contained (in mM): K-gluconate, 105; NaCl, 10; K_3_-citrate, 20; HEPES, 10; EGTA, 0.25; MgCl_2_, 0.5; Mg-ATP, 3; Na-GTP, 0.5. In most experiments, the pipette solution also included 2 µM Atto488. Recordings were digitized at 10-20 kHz and filtered with a Bessel filter at 2 kHz. For electrical stimulation of aPC projections, a glass pipette (resistance 1-2.5 MΩ), filled with ACSF and 2 µM Atto488, was located at the apical dendrite of the recorded granule cell with a distance of at least 50 µm. “Sniff cycle stimulation” was applied to evoke synaptic excitation of granule cells.

### Reagents

The reagents 2-Amino-5-phosphopentanoic acid (D-AP5, antagonist of NMDA-receptors), 2,3-Dioxo-6-nitro-1,2,3,4-tetrahydrobenzo[f]quinoxaline-7-sulfonamide (NBQX, antagonist of AMPA/kainate receptors), 4-[6-imino-3-(4-methoxyphenyl)pyridazin-1-yl]butanoic acid (Gabazine, antagonist of GABAA receptors) and N,2,2,3-tetramethylbicyclo[2.2.1]heptan-3-amine;hydrochloride (Mecamylamine hydrochloride, antagonist of nAChRs) were obtained from Alomone labs (Jerusalem, Israel). 2-Methyl-6-(phenylethynyl)pyridine hydrochloride (MPEP, antagonist of mGluR5-receptors), (*E*)-Ethyl 1,1a,7,7a-tetrahydro-7-(hydroxyimino)cyclopropa[*b*]chromene-1a-carboxylate (CPCCOEt, antagonist of mGluR1-receptors), 2’-Deoxy-*N*^6^-methyladenosine 3’,5’-bisphosphate ammonium salt (MRS2179, antagonist of P2Y_1_-receptors), (*R*)-(+)-7-Chloro-8-hydroxy-3-methyl-1-phenyl-2,3,4,5-tetrahydro-1*H*-3-benzazepine hydrochloride (SCH23390, antagonist of D1-receptors), [4-(4-Amino-6,7-dimethoxy-2-quinazolinyl)-1-piperazinyl]-2-furanylmethanone hydrochloride (Prazosin, antagonist of α1-adrenoreceptors), (α*S*)-α-(Hydroxymethyl)benzeneacetic acid (1α,2β,4β,5α,7β)-9-methyl-3-oxa-9-azatricyclo[3.3.1.0^2,4^]non-7-yl ester hydrobromide (Scopolamine, antagonist of mACh receptors) were purchased by Abcam (Cambridge, UK). The antagonists (2*S*)-3-[[(1*S*)-1-(3,4-Dichlorophenyl)ethyl]amino-2-hydroxypropyl](phenylmethyl)phosphinic acid hydrochloride (CPG55845, antagonist of GABAB-receptors), 17α-Hydroxy-20α-yohimban-16β-carboxylic acid,methyl ester hydrochloride (Rauwolscine hydrochloride, antagonist of α2-adrenoreceptors), and (±)-*erythro*-(*S*\*,*S*\*)-1-[2,3-(Dihydro-7-methyl-1*H*-inden-4-yl)oxy]-3-[(1-methylethyl)amino]-2-butanol hydrochloride (ICI 118,551 hydrochloride, antagonist of β-adrenoreceptors) were obtained from Tocris Bioscience (Bristol, UK). (*S*)-5-Aminosulfonyl-*N*-[(1-ethyl-2-pyrrolidinyl)methyl]-2-methoxybenzamide (Sulpiride, antagonist of D2-receptors) and 4-(-2-[7-amino-2-[2-furyl][1,2,4]triazolo[2,3-a][1,3,5]triazin-5-yl-amino]ethyl)phenol (ZM241385, antagonist of A_2A_ receptors) were obtained from Sigma-Aldrich (Merck, Darmstadt, Germany).

## Acknowledgments

We thank M. Rothermel (Aachen, Germany), F. Schneider-Warme (Freiburg, Germany) and F. Kirchhoff (Homburg, Germany) for providing transgenic mice. We thank A.C. Rakete and M. Fink for technical assistance. M. Doengi (Bonn, Germany) provided preliminary data. Schematic images were prepared using Biorender.com. We gratefully acknowledge financial support by the Deutsche Forschungsgemeinschaft (Project number 528690369 and SFB1328 TP-A07, project number 335447717).

## Author contributions

Study design: AB and CL. Methodology: AB, KL, JB, JSP, DD, CEG. Experiments: AB, KL, JB, JSP, NR, KS, DD, DH. Data analysis: AB, KL, JB, JSP, NR, KS, DD, DH, CL. Writing and figure design: AB, JB, CL. All authors edited and approved the manuscript.

## Data availability statement

All data are provided by the authors upon reasonable request.

## Supplementary Material

**Figure S1:**
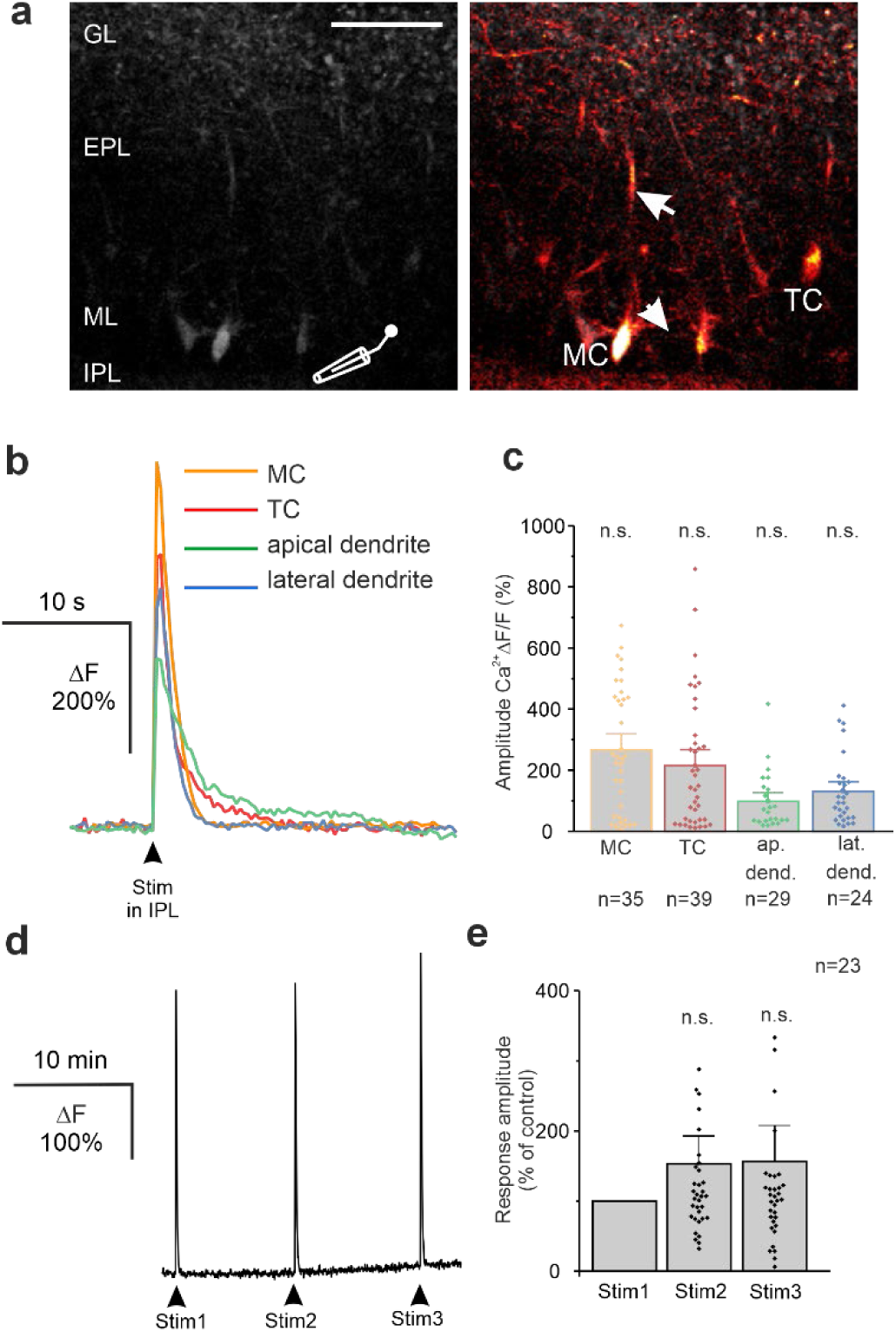
Stimulation-induced Ca^2+^ transients in mitral and tufted cells. **(a)**GCaMP6s-expressing mitral cells (MC) and tufted cells (TC) before (left) and during (right) electrical stimulation in the IPL. Ca^2+^ transients were recorded in somata, apical dendrites (arrow) and lateral dendrites (arrowheads). Scale bar: 50 µm. **(b)** Traces of GCaMP6s fluorescence changes upon electrical stimulation in the IPL. **(c)** The amplitudes of Ca^2+^ changes in MC, TC, apical dendrites (ap. dend.) and lateral dendrites (lat. dend.) were not significantly different. Since dendrites often could not be assigned to a mitral or tufted cell, we did not distinguish between dendrites of these two cell types. **(d)** Repetitive IPL stimulation did not lead to a significant decrease in the response amplitude in synaptic isolation established by glutamatergic (D-APV, NBQX, MPEP, CPCCOEt) and GABAergic (Gabazine, CGP55845) receptor antagonists in MCs. **(e)** Response amplitude upon repetitive stimulation in the IPL in MC normalized to the amplitude of the first stimulation. n.s., not significant.

**Figure S2:**
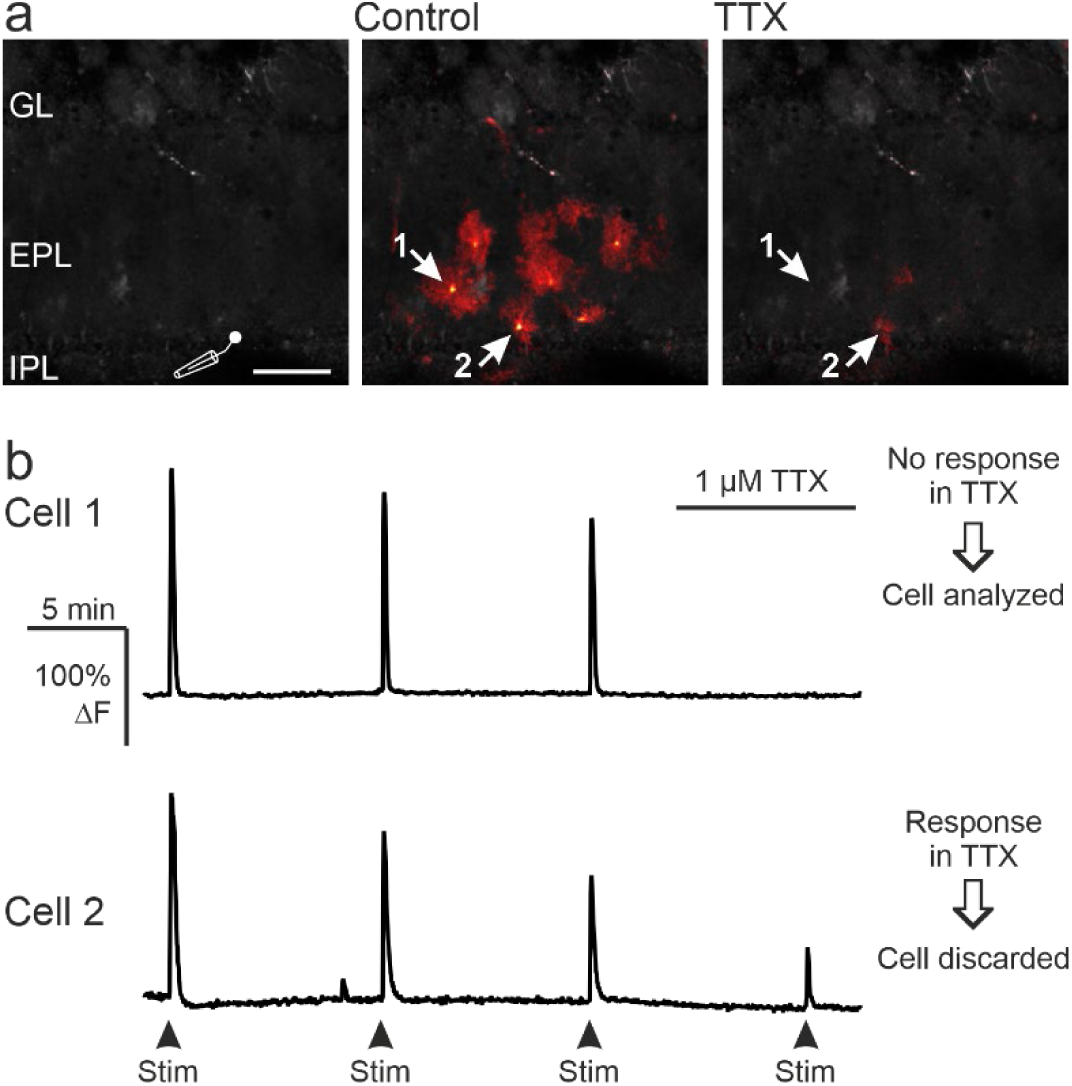
Effect of tetrodotoxin (TTX) on stimulation-induced Ca^2+^ transients in astrocytes. **(a)** Ca^2+^ transients evoked by electrical stimulation in the IPL in the absence (Control) and presence of TTX. Most astrocytes that responded to stimulation under control conditions did not respond in the presence of TTX (e.g. cell 1), whereas few astrocytes still generated small Ca^2+^ responses (cell 2). Scale bar: 50 µm. **(b)** Only astrocytes whose Ca^2+^ responses were entirely suppressed by TTX were considered for analysis (Cell 1), while astrocytes with residual Ca^2+^ response were discarded (Cell 2).

**Figure S3:**
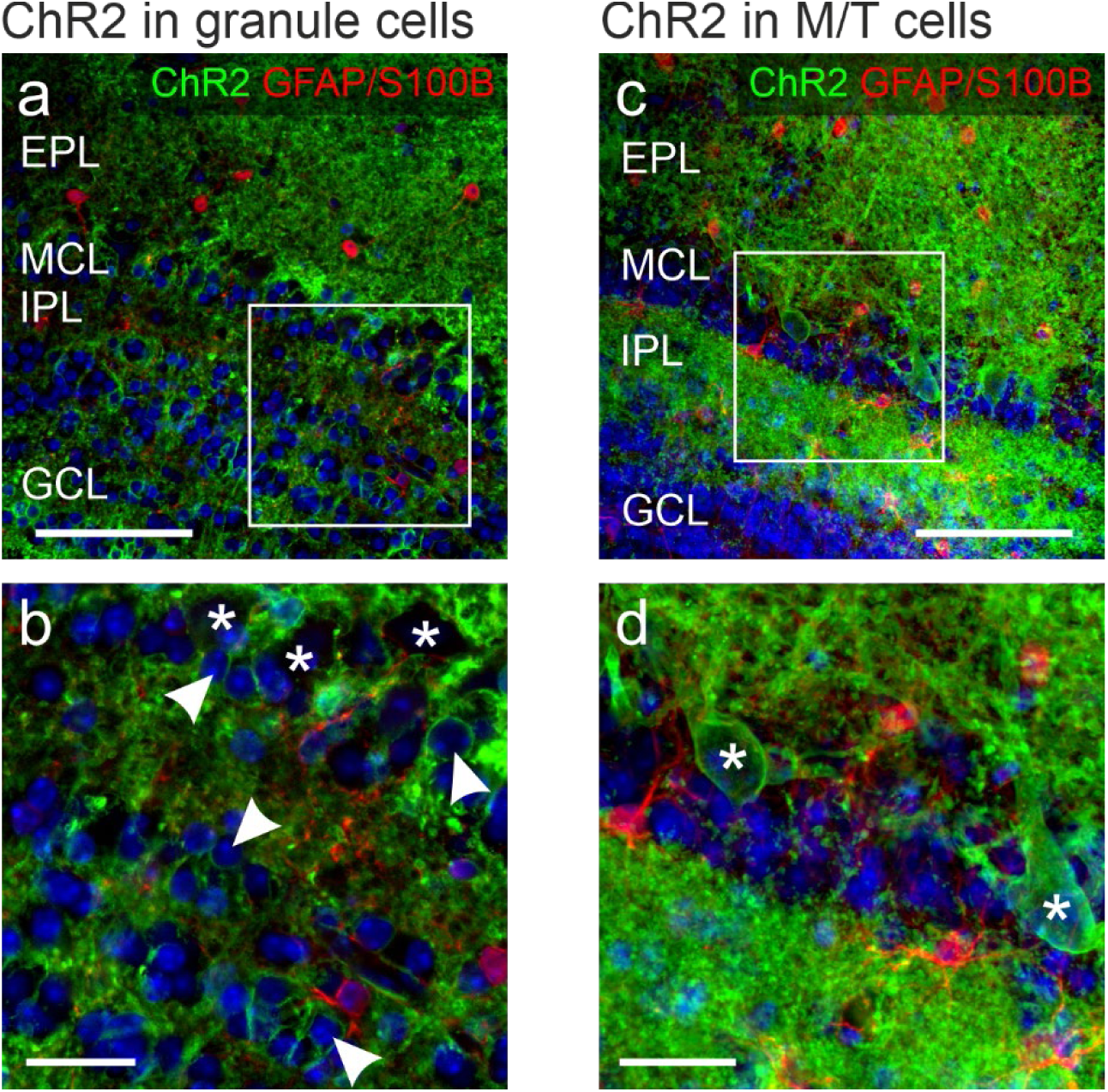
Expression of channelrhodopsin-2 in Pcdh21-Cre x ChR2^flox^ mice. **(a)** Immunohistochemistry of Pcdh21-Cre x ChR2^flox^ mice with ectopic expression pattern. ChR2^flox^ mice express EYFP which was visualized by an anti-GFP antibody (Green). **(b)** Magnification from (a). Ectopic expression of ChR2 in interneurons included GCs (arrowheads), but not astrocytes (red; immunolabeled with a combination of antibodies against GFAP and S100B) and MCs, as evident by the lack of ChR2-positive large cell bodies in the MCL (asterisks). Nuclei were stained with Dapi (blue). **(c)** In mice with entopic ChR2 expression, ChR2 was found in MCs, while no colocalization with astrocytes (red) and GCL (see granule cell layer; GCL) was detected. **(d)** Magnification from (c). Asterisks highlight ChR2/EYFP-positive MC cell bodies. Scale bars: a, c, 50 µm; b, d, 20 µm.

